# Exploring the Landscape of Spatial Transcriptome Analysis: Introducing STASH, a Database of Spatial Transcriptome Tools

**DOI:** 10.1101/2023.04.20.537419

**Authors:** Han Chu, Kun Wang, Hansen Cheng, Wenhao Ma, Liting Dong, Yixiong Gou, Jian Yang, Haoyang Cai

## Abstract

Spatial transcriptomics (ST) has emerged as a powerful tool for unravelling tissue structure and function. However, the continuous development of ST has made it challenging to select and effectively use appropriate analysis tools. To address this issue, we have developed the Spatial Transcriptome Analysis Hub (STASH, http://cailab.labshare.cn:7004), a comprehensive, systematic, and user-friendly database of ST analysis tools. STASH collects and categorizes most of the tools currently available and provides insight into their current status and trends. This can help researchers quickly locate the appropriate tool for their needs, or even guide researchers in the development of better tools.

## Introduction

In recent years, a number of ST technologies have emerged, enabling transcriptome analysis at the multi-cellular, single-cellular, or sub-cellular level while preserving spatial location information[1, 2]. These technologies can be broadly classified into two types: next-generation sequencing-based spatial transcriptomes (sST) and image-based spatial transcriptomes (iST)[3-5]. While sSTs allow whole transcriptome analysis, conventional methods such as 10X Visium[6] and slide-seq[7] are impeded by limited spatial resolution and low mRNA capture rates. Conversely, iSTs such as MERFISH[8] and seqFISH[9] offer high spatial resolution and mRNA capture rates but are restricted by the number of target genes and field of view sizes. Recently developed technologies, including HDST[10] and stereo-seq[11], are expected to offer alternatives with improved capabilities to overcome these limitations.

ST technologies have captured considerable attention in the research community, especially after being named Method of the Year 2020 by Nature Methods[12]. The bioinformatics community has developed a plethora of tools to analyze this new type of data with a spatial dimension. However, the lack of organization and categorization of these tools has made it challenging for researchers to develop and use ST analysis tools effectively. To address this issue, we have developed STASH (http://cailab.labshare.cn:7004), which collects and organizes the vast majority of tools available as of February 2023 and categorizes them based on ST data analysis tasks. In addition, by analyzing the database, we present trends related to each category of tools, including their development and application platforms, as well as the applicable ST technology types. We aim to facilitate researchers’ development and application by showing the current state of this rapidly evolving field.

## Construction and content

### Database

STASH serves as a comprehensive database of software tools specifically designed for the analysis of ST data. The collection of tools encompasses packages from repositories and code-sharing sites such as GitHub. In particular, the database provides important information such as DOIs, publication dates, and citation counts for publications related to the tools, including preprints. Each tool is accompanied by a brief description, its code repository, and relevant details such as the platform on which it was developed and used, as well as the applicable license.

To further streamline the tool selection process, we have categorized each tool according to the ST technologies to which it applies, based on technical documentation and descriptions in publications. In addition, we have categorized each tool according to the analysis tasks it can perform, allowing a more precise selection of tools for specific applications.

### Website

STASH, an online platform for spatial transcriptome analysis tools, was implemented by Golang (version 2.4.9), HTML, and JavaScript. Data storage is managed by MongoDB (version 2.4.9). The website has three main pages: ‘Search’, ‘Browse’, and ‘Analysis’. The ‘Browse’ page allows users to sort, retrieve and download tools based on various criteria. The ‘Search’ page provides detailed information about each tool, including descriptions, publication details, software code and license information, and associated software categories. The ‘Analysis’ page offers users an insight into current trends in spatial transcriptome research and provides an overview of the specific capabilities of each tool. The ‘Download’ page contains links to download the full information about STASH and the ‘Help’ page provides guidance on how to use the website.

### Analysis

The methodology employed in this study was based on the latest data that was available in STASH as of February 2023. Data processing was carried out using the R programming language (version 4.0.5) and the tidyverse package (version 1.1.0), while the resulting visualizations were generated using ggplot2 (version 3.3.3).

## Utility and discussion

### Overview of STASH

STASH is hosted on the Alibaba Cloud server (Ubuntu 14.04.2 LTS, GNU/Linux 3.13.0-32-generic i686), implemented by GO and MongoDB, and contains 260 tools (**Figure 1A**). At the outset of the database, we recorded 46 tools for ST analysis, which included most of the work up to December 2020. However, as of February 2023, we have incorporated 212 additional tools for ST analysis, representing a fivefold surge (**Figure 1B**). This remarkable increase in the number of tools reflects the growing attention of researchers to ST, along with its enhanced visibility and mounting commercialization (e.g., 10x Visium, Vizgen MERSCOPE, and NanoString CosMX).

**Figure 1.**
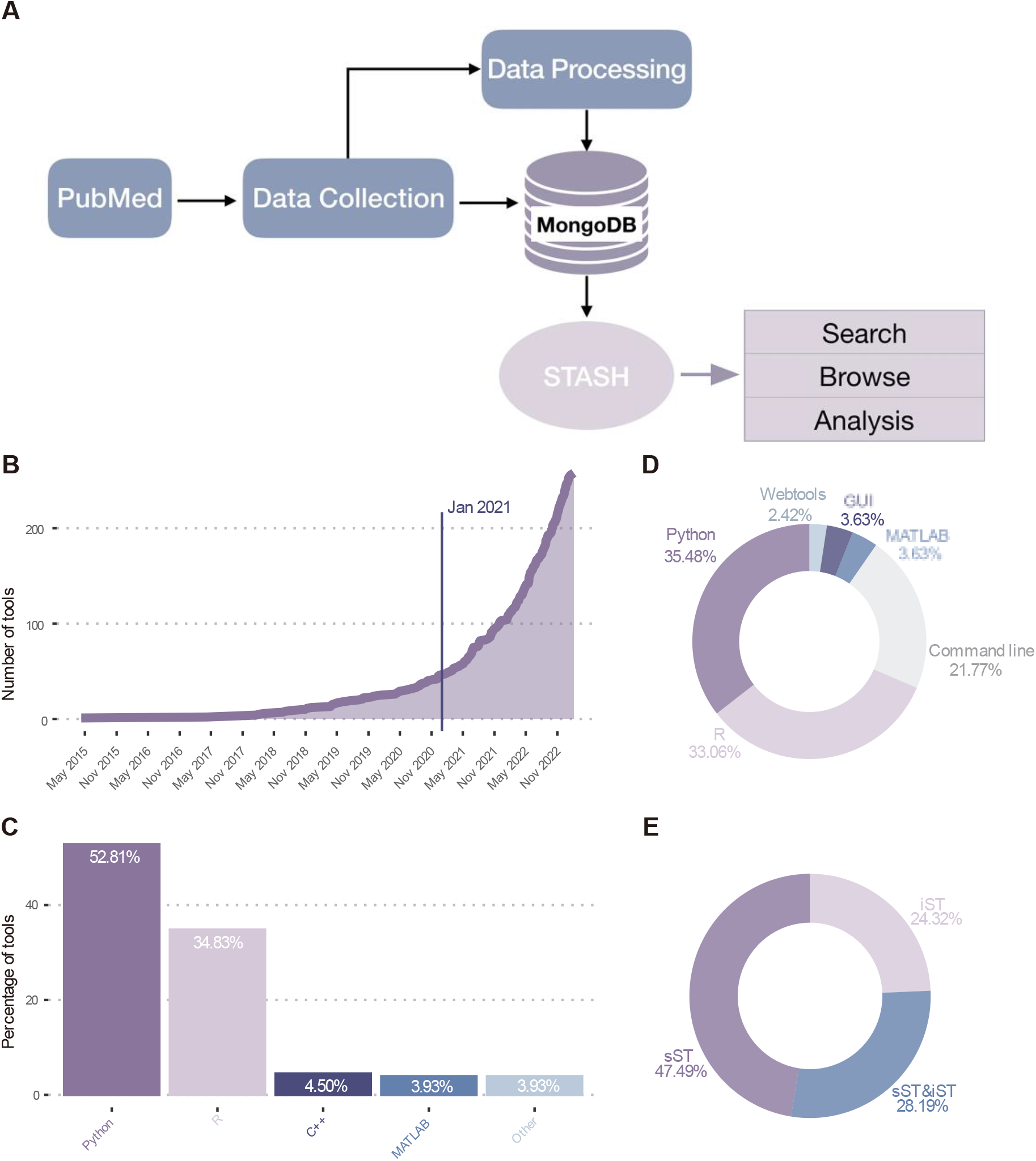
Overview of STASH. (A) A database design model that extracts key information from the articles collected in PubMed, then processes and stores the data in mongoDB, and finally makes it available for browsing on the website. (B) The number of tools for ST analysis increases over time, with a sharp increase after January 2021. (C) Platforms for the development of ST analysis tools. (D) Platform for the deployment of ST analysis tools. (E) Type of ST technology to which the ST tool applies.

#### Development Platform

To provide researchers with a better understanding of the current state of ST analysis tool development, we analyzed the existing development platforms (**Figure 1C**). Our statistical analysis shows that Python is currently the most widely used development tool. Its ability to handle histological image information and spatial location information contained in ST, as well as its suitability for deep learning, make it an excellent platform[13]. The statistical programming language R is the second most popular language after Python, with MATLAB and C++ in third and fourth place respectively. Notably, R has a centralized infrastructure, SpatialExperiment[14], designed specifically for the ST data format, which facilitates collaboration between small, specialized packages.

#### Application Platform

In terms of the platform used to analyze ST data, our analysis revealed that R and Python are the most prevalent, with 65% of available tools being R packages or Python modules (**Figure 1D**). Command line tools also show a growing trend in popularity (21%), giving data analysts greater flexibility to integrate results into their own analytical pipelines. Furthermore, a small number of graphical user interface tools and web-based tools offer a more user-friendly experience. Notably, transformations between the ST data format and R or Python platforms are already available, as demonstrated by the scDIOR[15]. It should be noted that the choice of platform may depend on the specific needs of the analysis and the expertise of the user.

#### Applicable ST technology types

To simplify the selection process of sST analysis tools, we have classified them according to the type of sST technology (**Figure 1E)**. Our results indicate that a greater number of analysis tools apply to sST compared to iST. This could be due to the resemblance between sST and widely used single-cell RNA sequencing (sc-RNA seq) techniques. However, it is important to recognize that different ST techniques may possess unique technical specifications and characteristics, even within the same type of ST technology. Therefore, data analysts must carefully evaluate the compatibility of a tool with the specific ST technique being used prior to implementation.

### Categories of ST analysis

ST technology has shown great potential in the analysis of gene expression profiles in tissues and is well suited to elucidate the cellular composition of tissues and the spatial interaction between different cell types, including molecular interactions between tissues[16]. In line with this analytical trend, the tools in STASH have been categorized into four analytical phases based on their function, including raw data processing, pre-processing, generic downstream tasks, and other downstream tasks (**Figure 2**).

**Figure 2.**
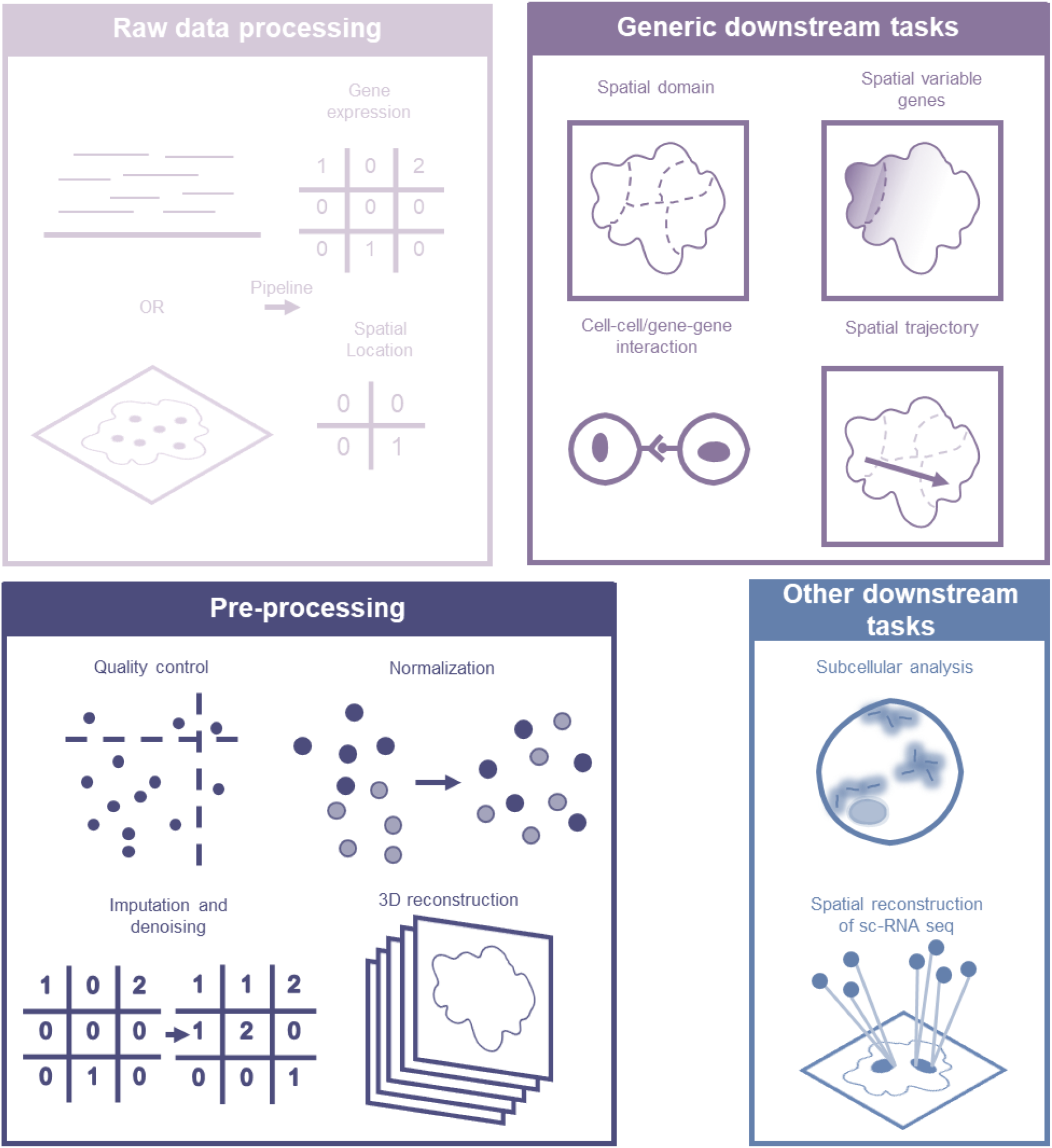
Phases of the ST analysis process based on current analysis trends. The processing of raw ST data allows the acquisition of gene expression profiles together with spatial location information. In order to obtain high quality data for downstream analysis, various methods are used in the pre-processing phase. Spatial domains and spatially variable genes constitute the primary focus of the general downstream analysis. Moreover, other downstream analyses extend the scope of ST data observations.

#### Raw data processing

The two primary types of ST technology, iST, and sST generate data through distinct processes. iST obtains gene expression matrix and spatial location information from microscopy images, requiring several steps such as spot detection, transcript decoding, cell segmentation, and transcript assignment. In contrast, sST acquires gene expression matrix and spatial location information from sequenced nucleotide sequences, with some steps in common with scRNA-seq, but also requires additional steps such as bead barcode decoding and spot and image alignment. Raw data processing pipelines (e.g., ST Pipeline [17] and STtools [18]) have been developed to process data from different ST technologies. Moreover, researchers have optimized specific steps of the pipeline, including cell segmentation. For most iSTs (as well as parts of sSTs), cell segmentation tools such as Baysor [19] and Cellpose[20] are useful to overcome challenges such as cell overlap and to enable analysis at single cell or even subcellular resolution.

#### Pre-processing

After obtaining the gene expression matrix and spatial location information, several pre-processing steps are required to ensure the accuracy of the downstream analysis. Initially, quality control is performed to remove unqualified cells or genes. Like scRNA-seq, ST would introduce unwanted technical artifacts that can be removed by normalization. However, it is important to note that Saiselet et al. have shown that ST data may not necessarily require normalization and that the total number of reads may be biologically relevant in terms of morphology and local cell density[21]. The limited number of target genes for iST and the low capture rate of mRNA in sST are accompanied by technical noise. To overcome these drawbacks, several gene imputation and denoising tools have been developed. An attractive aspect of ST is the integration of multiple samples to construct a stacked 3D alignment of the tissue, as exemplified by PASTE[22]. It is worth noting that ST integration is not limited to integration between slices, but its integration with other omics such as scRNA-seq (e.g., Seurat enables such integration[23]) can also provide new research perspectives. Imputation and integration are not necessarily mandatory for pre-processing. Dimensionality reduction tools that incorporate additional spatial location and tissue image information are categorized as spatial dimensionality reduction. Such tools, including SpatialPCA[24], can serve as a starting point for downstream analysis of ST data.

#### Generic downstream tasks

A high-quality expression matrix and associated features are obtained after pre-processing for downstream analysis. The identification of spatially coherent domains is crucial for the understanding of anatomical regions and requires the consideration of spatial information. Spatial dimensionality reduction can be used as a feature extraction during pre-processing to identify domains. If not, spatial domain identification tools such as BayesSpace[25] can be used to cluster cells based on spatial information. Genes with spatial expression patterns are critical determinants of polarity and anatomical structures, which can be identified using tools such as Spark[26]. For single-cell resolution ST, cell type identification can be achieved using a scRNA-seq reference or by examining markers of spatial domains. For multi-cellular resolution ST, spatial decomposition can be used to resolve the cell composition of each spot and identify the tissue cell composition. Once the cell type/composition is identified, the results of cell-cell/gene-gene interaction analysis (such as NCEM[27] have clear biological meaning. Similarly, spatial trajectory analysis can also provide valuable insights.

#### Other downstream tasks

Several tools are available to perform extended analyses on ST data beyond the general downstream tasks (Table 1). With high-resolution ST technologies, it is possible to perform subcellular analyses, such as spatial and temporal identification and analysis of transcripts within single cells using tools such as FISH-quant[28]. Another exciting area is the spatial reconstruction of scRNA-seq data, where ST data can be used as a reference to assess the spatial location of cells, as demonstrated by Tangram[4]. Also, several tools are being developed to detect biological signals other than expression in ST data, such as fusion transcripts[29], alternative polyadenylation[30], point mutation detection[31], CNV detection, and clone inference[32]. These tools expand the scope of what can be observed in ST data and offer new avenues for biological discovery.

**Table 1.** The division of the ST analysis phases and the description of categories for tools in STASH.

| Phase | Classification | Description |
| --- | --- | --- |
| Raw data processing | Bead barcode decoder | Bead barcode decoder of sST |
| Raw data processing | Cell segmentation | Identification of the precise boundaries of cells in images |
| Raw data processing | Image&spot alignment | Alignment of spot position in images |
| Raw data processing | Spot detection | Detection of transcript spot in image |
| Raw data processing | Nuclei Segmentation | Identification of cell nuclei in images |
| Raw data processing | Transcript decoder | Barcode decoder of iST |
| Raw data processing | Transcript assignment | Assignment of transcripts to cells |
| Raw data processing | Raw data processing pipeline | Comprehensive pipeline of raw data processing |
| Preprocessing | Quality control | Removal of low-quality cells (spots) and genes |
| Preprocessing | Imputation and denoise | Integration of sc-RNA data, spatial location, or histological image information for gene imputation or denoising |
| Preprocessing | Normalization | Removal of unnecessary changes affecting biological significance |
| Preprocessing | 3D reconstruction | Construction of stacked three-dimensional arrangement of tissues |
| Preprocessing | Slices alignment | Alignment and integration of slices |
| Preprocessing | Integration | Integration between ST data or with other omics data |
| Preprocessing | Spatial dimensionality reduction | Additional integration of spatial location information or histological images for dimensionality reduction |
| Generic downstream tasks | Super resolution | Profiling local gene expression patterns to improve ST data resolution |
| Generic downstream tasks | Cell type identification | Cell type identification of High-resolution ST |
| Generic downstream tasks | Spatial decomposition | Estimation of the proportion of cell types in spots |
| Generic downstream tasks | Spatial variable genes | Identification and analysis of genes with spatially variable correlated structures (density-related) |
| Generic downstream tasks | Gene co-expression | Gene co-expression analysis |
| Generic downstream tasks | Spatial expression modules | Identification and analysis of gene set with spatial pattern |
| Generic downstream tasks | Cell-cell/gene-gene interaction | Analysis of cell-to-cell interactions or receptor-ligand binding in tissues |
| Generic downstream tasks | Spatial domain | Cell (spot) clustering with expression profiles, spatial location information, or histological images |
| Generic downstream tasks | Cell co-location | Co-localization analysis of cells |
| Generic downstream tasks | Spatial trajectory | Ordering of cells along a trajectory with spatial location information |
| Other downstream tasks | Fusion transcript | Detection and analysis of fusion transcripts |
| Other downstream tasks | Spatial heterogeneity | Heterogeneity analysis of complex environments in spatial molecular biology |
| Other downstream tasks | Neuronal activity quantitation | Quantification of neuronal activity |
| Other downstream tasks | Gene panel selection | Gene panel selection for the target of iST |
| Other downstream tasks | Alternative Polyadenylation | Detection and analysis of Alternative Polyadenylation |
| Other downstream tasks | Subcellular analysis | Identification and analysis of spatial and temporal dynamics of transcripts within single cells |
| Other downstream tasks | Spatial reconstruction of SC | Spatial reconstruction of sc-RNA data |
| Other downstream tasks | CNV detection | Detection of copy number variation |
| Other downstream tasks | Clone inference | Inference of clonal relationships |
| Other downstream tasks | Point Mutation Detection | Detection of point mutation |
| Other downstream tasks | Allelic | Identification and analysis of Allelic |
| Multiple | Visualization | Visualization of spatial transcriptome data for multiple processes |
| Multiple | Interactive | Interactive components or graphical user interfaces tools |

### Trends in ST analysis tasks

Each of the tools integrated into STASH covers one or more categories of analysis, as shown in **Figure 3A**. While some tools provide multiple categories of analysis in the form of raw data processing pipelines or toolboxes, most are tailored to perform specific ST analysis tasks. Toolboxes such as Squidpy[33], SPATA[34], and Giotto[35] provide a wide range of analyses from the expression matrix, making them ideal for exploratory data analysis. However, it should be recognized that the functionality of these toolboxes may not always be up to date with the latest developments in the corresponding functions.

**Figure 3.**
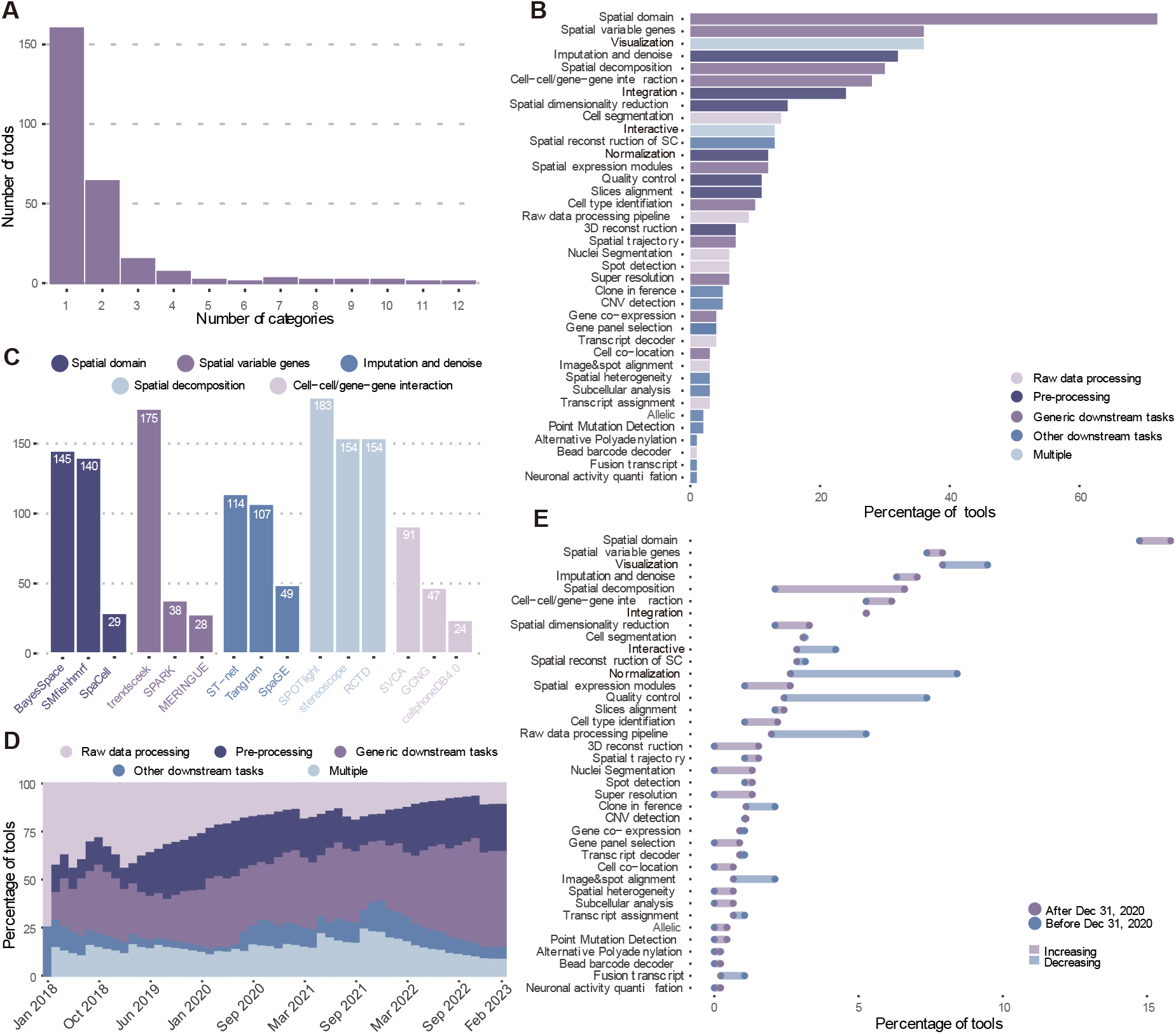
Trends in ST analysis tasks. (A) The number of categories associated with each tool in STASH. (B) The categories of ST analysis tools in STASH (a tool may contain more than one category). (C) The top three most cited non-toolbox tools in the highest-ranked categories. (D) The percentage of tools in different stages of ST varies over time. (E) Change in the percentage of ST analysis tools in each category before and after being named Method of the Year.

We conducted a comprehensive analysis of all analysis categories (**Figure 3B**) and presented the top three most cited non-toolbox tools in the highest-ranked categories (**Figure 3C**). The results reveal that the spatial domain is the most popular analysis category among researchers. Although combining spatial location information with histological image information would improve spatial domain identification, existing tools fail to provide the required level of stability, and further improvements in information extraction and feature selection are needed for both spatial and histological data[36]. The analysis also shows that spatially variable genes and visualization are the second and third most popular analysis categories. However, it is important to exercise caution when drawing biological conclusions based on genes obtained from statistical significance-based screening tools for spatially variable genes, regardless of the method used[37]. This highlights the need for further refinement of spatially variable gene tools.

We found that only imputation and denoising ranked highly in the pre-processing phase, highlighting the limitations of current ST technologies in terms of mRNA capture rate or the number of targeted genes. Various strategies currently exist for tools in this category, including scRNA-seq-based approaches such as iSpatial[38], tissue images information-based methods such as xfuse[39], spatial location information-based techniques such as DIST[40], and multiple information-based methods such as Sprod[41]. Spatial decomposition, a critical step in obtaining tissue cell composition using low-resolution ST, requires scRNA-seq data as a reference. RCTD is considered one of the better tools in this category, as several studies have shown[42, 43]. Although ranked sixth, cell-cell/gene-gene interaction is still attracting considerable interest from researchers, suggesting the possibility of future developments in this area.

The analysis showed that the proportion of tools for general downstream tasks has gradually increased over time, while the proportion of tools for raw data processing has decreased (**Figure 3D**). Using January 2021 as the cut-off point, we tracked changes in the proportions of different categories among all tools (**Figure 3E**). In particular, the raw data processing pipeline has decreased significantly, suggesting that ST technology is maturing and more attention is being directed toward downstream analysis. On the other hand, there has been a significant increase in spatial decomposition tools, suggesting a breakthrough in processing strategies for low-resolution ST, which has been a long-standing challenge for researchers in this field.

## Conclusions

Over the past few years, the number of software tools available for ST analysis has increased significantly, with all over 250 currently available. These tools have been cataloged and classified in STASH. Our analysis of the database has shown that researchers are predominantly interested in the generic downstream tasks, particularly those in the spatial domain. Although derived from different ST technologies, databases such as SODB provide data with a uniform data format and provide an excellent basis for the development of ST data analysis tools[44].

As ST technologies continue to advance, it is expected that new tools will emerge to improve ST data analysis or extend the scope of ST data observations. Tools that perform well, continue to develop, and provide an excellent user experience are expected to become the standard analysis choice as detailed benchmarks and comparisons are performed. High-resolution ST, including single cell and sub-cellular resolution, with a wide field of view size, may represent the future direction of technical progress and require the development of new tools. Moreover, the integration of ST with other omics technologies, such as scRNA-seq, represents another area of growth.

As the field advances, STASH will continue to receive updates from the community to facilitate the effective development and application of ST data analysis tools by researchers.

## Declarations

### Ethics approval and consent to participate

Not applicable

### Consent for publication

Not applicable

### Availability of data and materials

The datasets generated and/or analysed during the current study are available in the website, [http://cailab.labshare.cn:7004]

### Author contributions

H.C. conducted most experiments and analyzed the data. K.W. built the wedsite. H.C., K.W. and H.Y.C. designed all experiments. H.S.C., W.H.M., L.T.D, Y.X.G, and J.Y. helped with the collection of the public data. H.C. prepared the manuscript. H.Y.C. directed the research.

## Acknowledgements

Not applicable

